# Myogenetic Oligodeoxynucleotide Induces Myocardial Differentiation of Murine Pluripotent Stem Cells

**DOI:** 10.1101/2023.07.31.551374

**Authors:** Mina Ishioka, Yuma Nihashi, Yoichi Sunagawa, Koji Umezawa, Takeshi Shimosato, Hiroshi Kagami, Tatsuya Morimoto, Tomohide Takaya

## Abstract

An 18-base myogenetic oligodeoxynucleotide (myoDN), iSN04, acts an anti-nucleolin aptamer and induces myogenic differentiation of skeletal muscle myoblasts. This study investigated the effect of iSN04 on murine embryonic stem cells (ESCs) and induced pluripotent stem cells (iPSCs). In the undifferentiated state, iSN04 inhibited the proliferation of ESCs and iPSCs but did not affect the expression of pluripotent markers. In the differentiating condition, iSN04 treatment of ESCs/iPSCs from day 5 onward dramatically induced the differentiation into *Nkx2-5*^+^ beating cardiomyocytes with upregulation of *Gata4, Isl1*, and *Nkx2-5*, whereas iSN04 treatment from earlier stages completely inhibited cardiomyogenesis. RNA sequencing revealed that iSN04 treatment from day 5 onward contributes to the generation of cardiac progenitors by modulating the Wnt signaling pathway. Immunostaining showed that iSN04 suppressed the cytoplasmic translocation of nucleolin and restricted it to the nucleoli. These results demonstrate that nucleolin inhibition by iSN04 facilitates the terminal differentiation of cardiac mesoderm into cardiomyocytes, but interferes with the differentiation of early mesoderm into the cardiac lineage. This is the first report on the generation of cardiomyocytes from pluripotent stem cells using a DNA aptamer. Since iSN04 did not induce hypertrophic responses in primary-cultured cardiomyocytes, iSN04 would be useful and safe for the regenerative therapy of heart failure using stem cell-derived cardiomyocytes.

## 1. Introduction

Pluripotent stem cells (PSCs), such as embryonic stem cells (ESCs), have unlimited self-renewal and pluripotency to generate three germ layers that differentiate into all cell lineages, which can be used for regenerative therapy of various neurological, metabolic, and cardiovascular diseases [1]. Induced PSCs (iPSCs) exhibit self-renewal and pluripotency to the same extent as ESCs, which are typically generated by introducing transcription factors (Oct3/4, Sox2, Klf4, and c-Myc) into somatic cells [2,3]. Since the patient-derived iPSCs are immunologically suitable sources for cell transplantation into themselves, the technologies to direct iPSCs into specific cell lineages have been intensively studied. However, these protocols are often complicated and require expensive materials, including growth factors and basal matrix. Convenient and reproducible methods to differentiate iPSCs into the desired cell lineages are needed to make regenerative medicine feasible. Nucleic acid aptamers are single-stranded oligonucleotides that specifically bind to their target proteins in a conformation-dependent manner, similar to the antigen-antibody reaction. Aptamers will be favorable molecules to regulate stem cell fate in clinical settings, because they can be economically synthesized on a large scale, chemically modified for delivery, and thermally stable for storage [4]. For example, the assembly of DNA aptamers targeting the fibroblast growth factor (FGF) receptor can mimic basic FGF and support self-renewal of human iPSCs [5]. This suggests that aptamers have the potential to control the proliferation and differentiation of PSCs.

We have recently reported that a series of myogenetic oligodeoxynucleotides (myoDNs), which are 18-base telomeric DNAs designed from the lactic acid bacterium genome, facilitate the myogenic differentiation of skeletal muscle myoblasts and rhabdomyosarcomas [6-10]. One of the myoDNs, iSN04 (5’-AGA TTA GGG TGA GGG TGA-3’), acts as an anti-nucleolin aptamer and improves p53 protein levels by reversing nucleolin-inhibited translation of p53 mRNA, resulting in induction of myogenesis [6]. Nucleolin is a multifunctional phosphoprotein that is ubiquitously expressed and localized to the nucleus, cytoplasm, or plasma membrane depending on the context of cellular processes such as gene expression, protein shuttling, cytokinesis, and apoptosis [11]. The role of nucleolin in PSCs has been consecutively studied. Nucleolin is involved in the formation of nucleolus precursor bodies during early proliferation of porcine zygotes [12]. In murine ESCs (mESCs), downregulation of nucleolin impairs cell growth and survival [13]. Phosphorylated nucleolin physically interacts with the tumor protein Tpt1 during mitosis and with Oct4 during interphase in mESCs to regulate cell proliferation and differentiation [14]. Nucleolin also suppresses p53 protein levels and its downstream signaling pathway to maintain self-renewal of mESCs [15]. Nucleolin forms a complex with a retrotransposon LINE1, binds to ribosomal DNA to promote ribosomal RNA synthesis, and ultimately contributes to self-renewal of mESCs [16]. These reports indicate that nucleolin is an important for the proliferation of undifferentiated ESCs. However, the function of nucleolin during the differentiation of PSCs remains unclear.

This study investigated the effect of nucleolin inhibition by iSN04 on mESCs and murine iPSCs (miPSCs). Interestingly, iSN04 induced myocardial differentiation depending on the time of treatment, in part by modulating the Wnt signaling pathway and the intracellular localization of nucleolin. It indicates that iSN04 can be a useful aptamer for generating cardiomyocytes from PSCs.

## 2. Results

### 2.1. iSN04 Inhibits Proliferation of Undifferentiated PSCs

A miPSC line 20D17 expresses green fluorescent protein (GFP) under the control of the *Nanog* gene, a marker of undifferentiated PSCs [17]. Undifferentiated 20D17 cells were maintained on a feeder layer of mitomycin C (MMC)-treated murine embryonic fibroblasts (MEF) in growth medium (GM). The 20D17 cells were then treated with iSN04, and the *Nanog*-GFP^+^ colony size was quantified as an index of cell proliferation. As shown in Figure 1A, iSN04 significantly reduced the *Nanog*-GFP^+^ colony size without affecting *Nanog*-GFP expression. Quantitative real-time RT-PCR (qPCR) showed that iSN04 treatment did not alter the mRNA levels of the endogenous undifferentiation marker genes, *Klf4, Nanog, Pou5f1* (Oct4), and *Sox2* in 20D17 cells (Figure 1B). Similarly, in a mESC line hCGp7, iSN04 significantly reduced the size of colonies with enzymatic activity of alkaline phosphatase (ALP), another undifferentiation marker of PSCs (Figure 1C). These results demonstrate that iSN04, an anti-nucleolin aptamer, inhibits proliferation but does not induce differentiation of murine PSCs in the undifferentiated state, which is consistent with the previous studies reporting that downregulation of nucleolin impairs the growth of mESCs [13,15,16].

**Figure 1.**
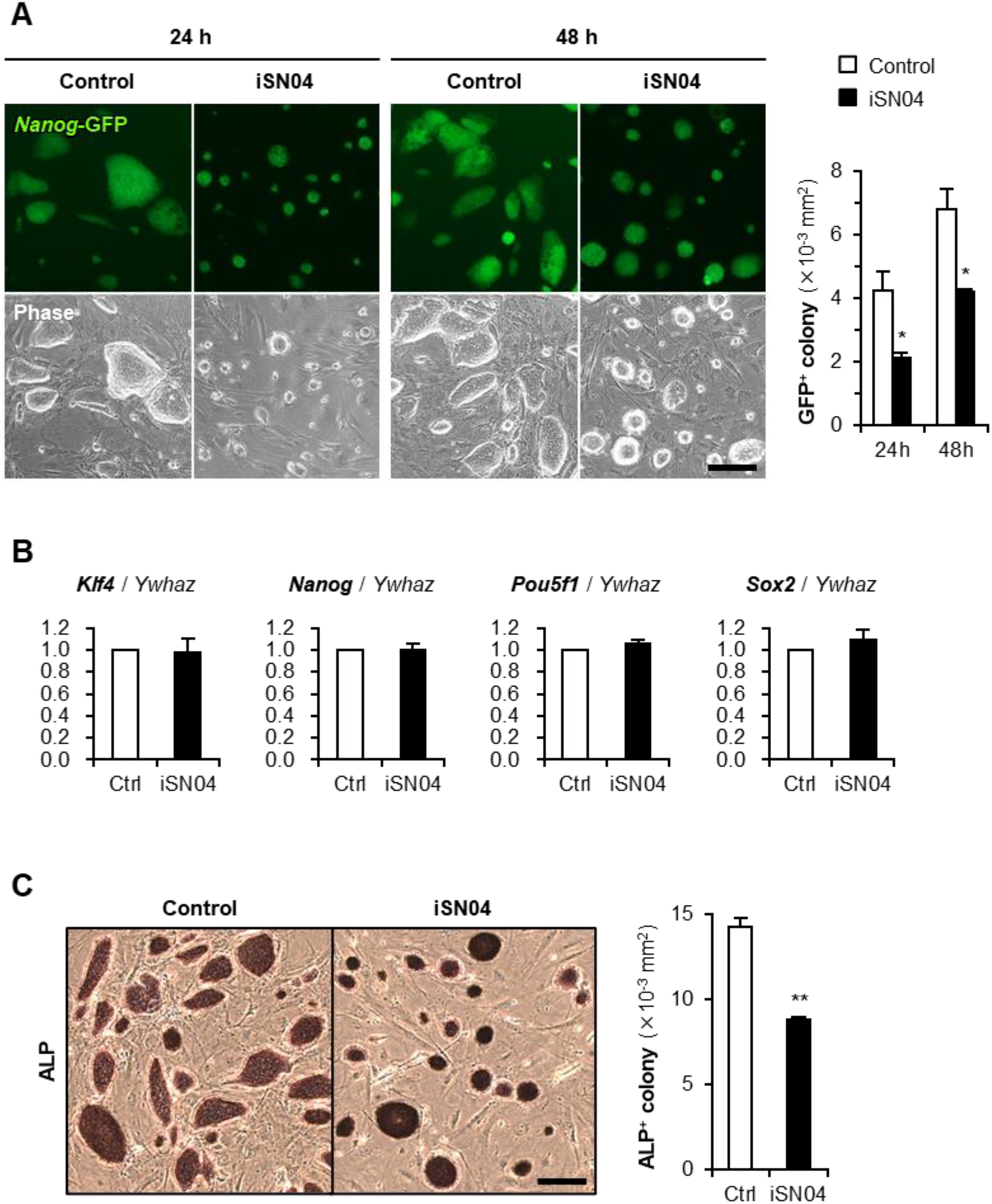
iSN04 inhibits proliferation of undifferentiated miPSCs and mESCs. (**A**) Representative fluorescence images of 20D17 cells treated with 10 μM iSN04 in GM for 24 and 48 h. Scale bar, 200 μm. *Nanog*-GFP^+^ colony size was quantified. * *p* < 0.05 vs control (Student’s *t*-test). *n* = 4 fields. (**B**) qPCR results of 20D17 cells treated with 10 μM iSN04 in GM for 48 h. *n* = 3. (**C**) Representative images of ALP staining of hCGp7 cells treated with 10 μM iSN04 in GM for 48 h. Scale bar, 200 μm. ALP^+^ colony size was quantified. ** *p* < 0.01 vs control (Student’s *t*-test). *n* = 4 fields.

### 2.2. iSN04 Induces Myocardial Differentiation of PSCs

The effect of iSN04 on the spontaneous differentiation of murine PSCs was investigated. 20D17 cells were induced to differentiate in differentiation medium (DM) and treated with iSN04 from day 5 to day 9. At day 11, a few beating clusters per dish (∼300 μm diameter and ≈50 beats/min), which appeared to be cardiomyocytes, were observed in the control group (Supplementary Video S1). Interestingly, in the iSN04-treated group, the large and vigorously beating clusters (>1 mm diameter and >100 beats/min) were obtained throughout the dishes (Supplementary Video S2). This suggests that iSN04 facilitates myocardial differentiation of PSCs.

To confirm whether the iSN04-induced beating cells were cardiomyocytes, hCGp7 cells expressing GFP under the control of the *Nkx2-5* gene [18,19], one of the earliest cardiac markers, were induced to differentiate in DM and treated with iSN04 from day 5 on 30-mm dishes. As shown in Figure 2A, *Nkx2-5*-GFP^+^ beating clusters were observed in the iSN04-treated group but not in the control group at day 8, indicating that the iSN04-induced beating clusters were undoubtedly cardiomyocytes.

**Figure 2.**
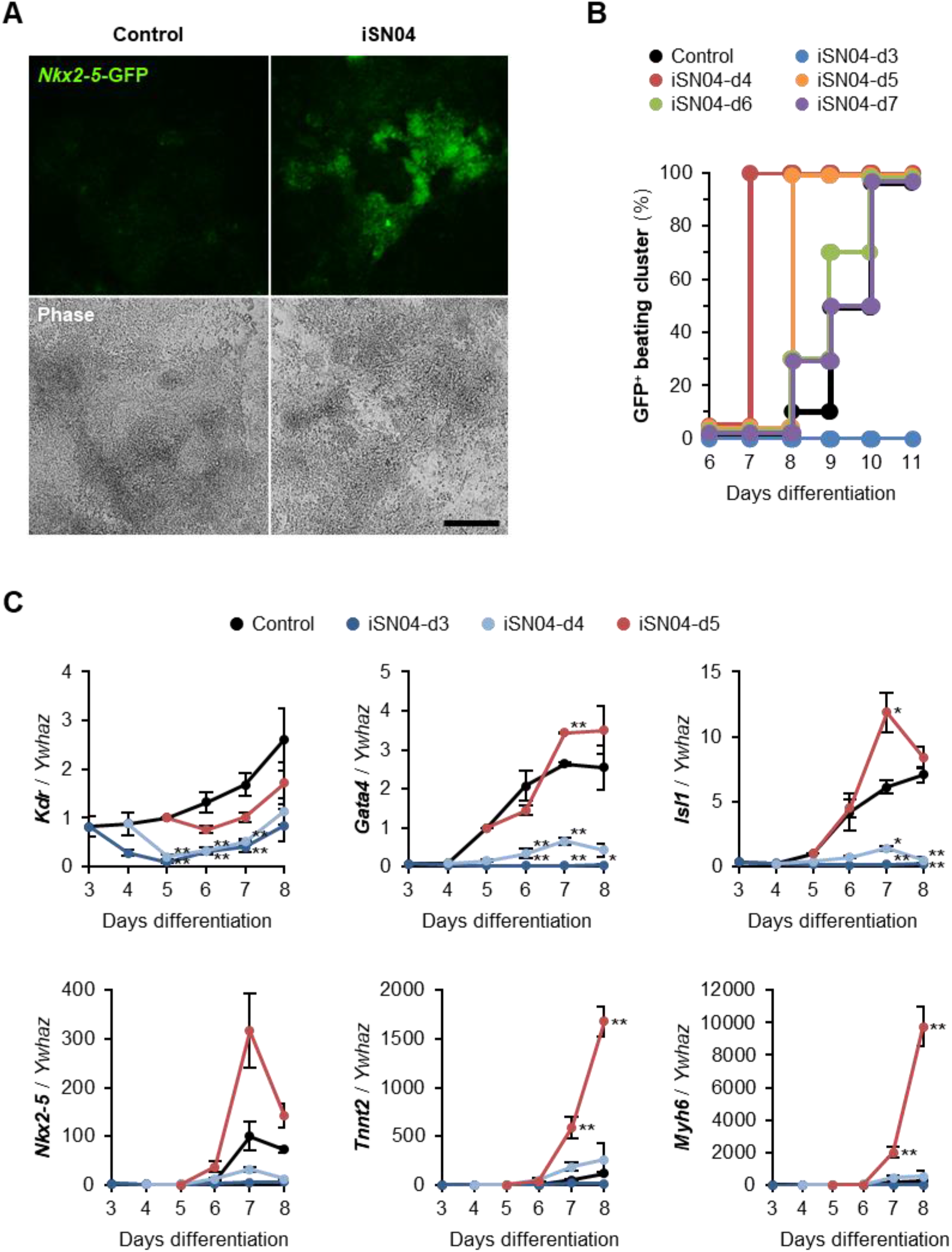
iSN04 induces myocardial differentiation of mESCs. (**A**) Representative fluorescence images of hCGp7 cells induced to differentiate in DM and treated with 10 μM iSN04 from day 5 to day 8 on 30-mm dishes. Scale bar, 200 μm. (**B**) hCGp7 cells induced to differentiate in DM and treated with 10 μM iSN04 from day 3, 4, 5, 6, or 7 on 96-well plates. Cumulative percentages of wells in which *Nkx2-5*-GFP^+^ beating clusters were observed are shown. *n* = 10. (**C**) qPCR results of hCGp7 cells induced to differentiate in DM and treated with 10 μM iSN04 from day 3, 4, or 5 on 30-mm dishes. * *p* < 0.05, ** *p* < 0.01 vs control on each day (Tukey-Kramer test). *n* = 3.

To investigate the time-dependence of the effect of iSN04 during differentiation, hCGp7 cells were seeded on 96-well plates and treated with iSN04 from day 3, 4, 5, 6, or 7 (iSN04-d3 – d7). As shown in Figure 2B, GFP^+^ beating clusters were observed in the control group from day 8 and appeared in all wells at day 10. iSN04-d6 and -d7 slightly increased the production of GFP^+^ myocardial clusters. Furthermore, iSN04-d4 and -d5 significantly accelerated myocardial differentiation, with GFP^+^ clusters reaching 100% at day 7 and 8, respectively. However, iSN04-d3 completely inhibited differentiation into *Nkx2-5*-GFP^+^ cells at least until day 11. These data demonstrate that the iSN04 switches the fate of PSCs into cardiac or non-cardiac lineage depending on the stage of differentiation.

The time-dependent effect of iSN04 on cardiac gene expression was examined by qPCR. hCGp7 cells were seeded on 30-mm dishes, induced to differentiate in DM, and treated with iSN04 from day 3, 4, or 5. In this experiment, iSN04-d3 and -d4 completely inhibited but iSN04-d5 significantly induced cardiomyogenesis. The result of iSN04-d4 on 30-mm dishes was different from the previous experiment on 96-well plates (Figure 2B), probably due to the subtle difference in culture conditions between the 30-mm dishes and the 96-well plates. It strongly suggests that the differentiation period around day 4 is a key stage for iSN04-driven cardiac lineage production. As shown in Figure 2C, iSN04-d3 and -d4 significantly reduced the mRNA levels of *Kdr* (Flk1; vascular endothelial growth factor receptor 2), a marker of mesoderm including prospective cardiovascular progenitors. The expression patterns of *Gata4* (a transcription factor for heart development), *Isl1* (a cardiac progenitor marker), and *Nkx2-5* were analogous; iSN04-d3 and -d4 significantly suppressed their expression throughout differentiation, whereas iSN04-d5 strongly induced them at day 7. As a result, the mRNA levels of *Tnnt2* (cardiac troponin T) and *Myh6* (cardiac α-myosin heavy chain), which are sarcomeric proteins expressed in the terminally differentiated cardiomyocytes, were massively increased in the iSN04-d5 group. These results suggest that administration of iSN04 at earlier stages disrupts mesodermal differentiation into the cardiac lineage, resulting in poor generation of cardiac progenitors. On the other hand, stimulation of the late mesoderm or cardiac mesoderm by iSN04 activates the myocardial program to produce mature cardiomyocytes.

### 2.3. iSN04 Influences Mesoderm Differentiation into Cardiac Progenitors

For a comprehensive analysis of iSN04-dependent gene expression in hCGp7 cells, total RNAs from the control, iSN04-d4, and iSN04-d5 groups at days 4-7 used for the qPCR (Figure 2C) were subjected to RNA sequencing (RNA-seq). 54.8 million (M) raw reads (average per sample) were processed into 54.6 M clean reads, of which 50.0 M reads (91.2%) were mapped to the mouse reference genome (Supplementary Table S2). Gene expression levels were calculated as the fragments per kilobase per million reads (FPKM). As shown in Supplementary Figure S1, the FPKM values defined by RNA-seq were highly correlated with the qPCR results, indicating that the RNA-seq data represent gene expression patterns in the iSN04-treated hCGp7 cells. The 13,613 genes expressed as FPKM > 1.0 in any sample were subjected to the following analysis.

To investigate the effect of iSN04 on the global gene expression patterns, principal component analysis (PCA) was performed using the FPKM values of all the 9 samples defined by RNA-seq. PCA identified the first, second, and third principal components (PC1, PC2, and PC3) with contributions of 0.52, 0.26, and 0.09, respectively. The 9 samples were plotted on the PCA spaces reconstructed by PC1, PC2, and PC3 (Supplementary Figure S2). However, there is no obvious relationship between the principal components and iSN04 treatments, suggesting that the impact of iSN04 on the transcriptome of differentiating PSCs is limited.

According to the qPCR results, the FPKM values of individual mesodermal and cardiac marker genes were surveyed. As shown in Figure 3, the expression levels of the precardiac mesoderm markers *T* (brachyury), *Mesp1, Mesp2*, and *Msgn1* (mesogenin 1) were peaked at day 6. Their transcriptions were downregulated in both iSN04-d4 and -d5 groups. The levels of cardiac mesoderm markers *Kdr, Pdgfra* (platelet-derived growth factor receptor α), and *Gata4* were downregulated in the iSN04-d4 group but not significantly altered in the iSN04-d5 group. The expression of *Mef2a*, a transcription factor required for cardiac reprogramming with *Gata4* [20], was not altered at all in either iSN04-d4 or -d5 group. The expression levels of the cardiac progenitor markers *Isl1, Nkx2-5, Tbx5* (a transcription factor cooperating with *Nkx2-5*), and *Hand2* (a transcription factor in the second heart field progenitors) at day 7 were apparently increased in the iSN04-d5 group, whereas those of *Isl1* and *Hand2* were decreased in the iSN04-d4 group. As a result, the sarcomeric genes *Actc1* (cardiac α-actin), *Tnnt2, Myh6*, and *Myl7* (myosin light chain 7) were strikingly induced in the iSN04-d5 group. These data suggest that iSN04 treatment of cardiac mesoderm facilitates differentiation into cardiac progenitors, leading to terminal differentiation into cardiomyocytes.

**Figure 3.**
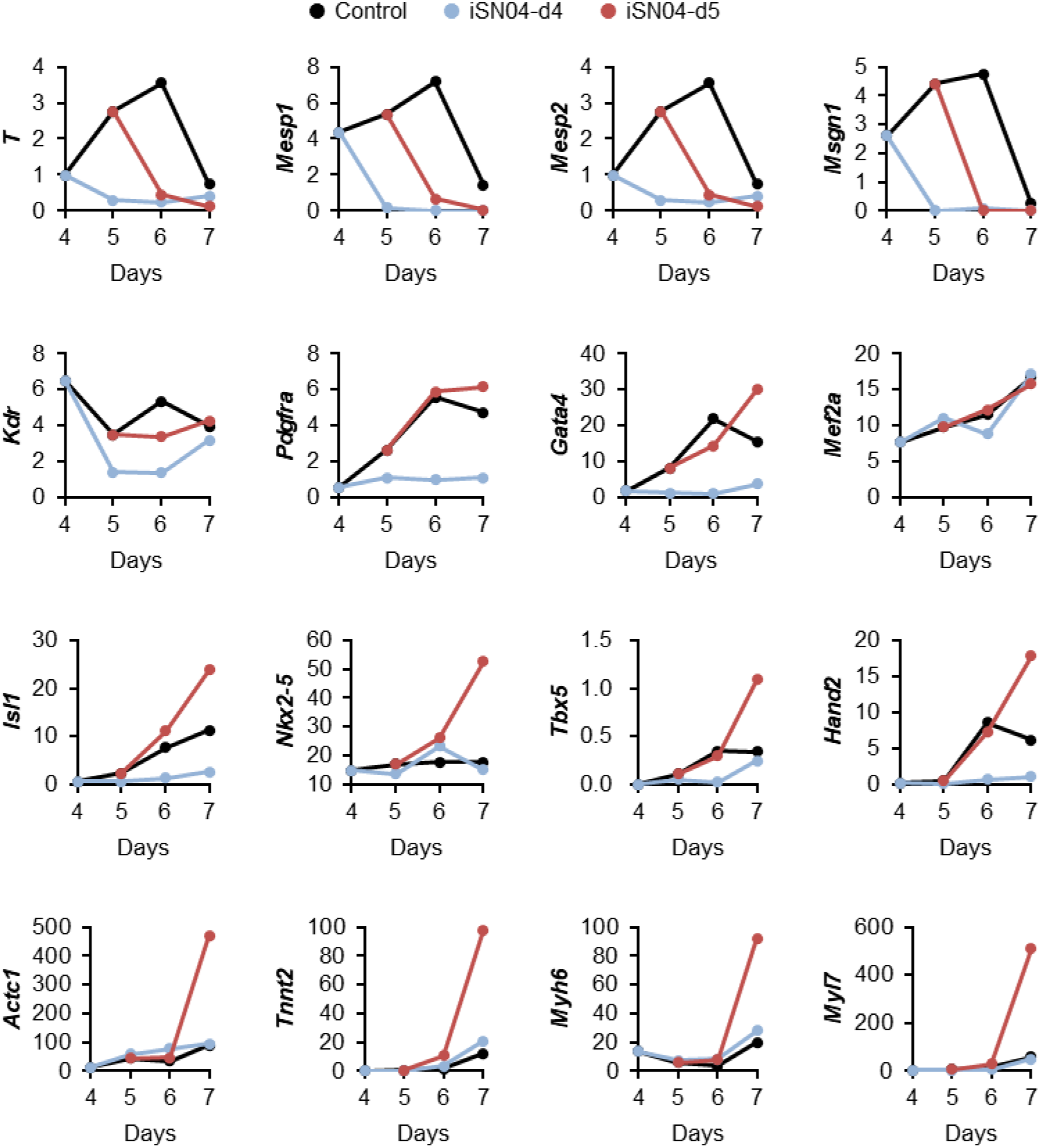
iSN04-dependent expression of mesodermal and cardiac genes in mESCs. RNA-seq results of hCGp7 cells induced to differentiate in DM and treated with 10 μM iSN04 from day 4 or 5 on 30-mm dishes (same samples as in Figure 2C). FPKM values are displayed as expression levels.

### 2.4. iSN04 Affects the Wnt Signaling Pathway in PSCs

From the FPKM values of the 13,613 genes, the differentially expressed genes (DEGs) among the samples were defined by considering |fold-change| ≥ 4.0 as a cutoff. The highly expressed DEGs in the control, iSN04-d4, and iSN04-d5 groups were designated as High-ctrl, High-d4, and High-d5, respectively (Figure 4). At day 6, the 606 DEGs included 287 High-ctrl, 163 High-d4, and 156 High-d5 genes. At day 7, the 455 DEGs involved 151 High-ctrl, 140 High-d4, and 164 High-d5 genes. These DEG subsets were subjected to KEGG pathway analysis. As listed in Table 1, the genes involved in the High-ctrl and High-d5 groups were significantly enriched in the Wnt signaling pathway (*p* < 0.05), whereas the High-d4 groups had no Wnt-related genes. The High-ctrl groups (i.e. down-regulated by iSN04) at day 6 and 7 contained 6 genes in common; *Cer1* (Cerberus), *Dkk1* (Dickkopf1), *Fzd10* (Frizzled homolog 10), *Sox17, Wnt3*, and *Wnt6*. Of these, *Cer1* and *Dkk1* are Wnt antagonists and inducers of cardiac mesoderm [21]. On the other hand, *Fzd10* is a Wnt receptor and *Wnt3/6* are Wnt ligands that activate the canonical Wnt/β-catenin pathway [22]. It has been reported that *Wnt3* induces mesodermal differentiation of PSCs [23], and *Wnt6* inhibition promotes cardiac progenitor differentiation [24]. *Sox17* induces endoderm and neuromesoderm differentiation of PSCs as downstream of β-catenin signaling [25]. These data demonstrate that iSN04 disrupts the Wnt/β-catenin signaling pathway in PSCs, which is known to play a stage-specific dual role during differentiation, directing early mesoderm toward the cardiac lineage but inhibiting terminal differentiation into cardiomyocytes at a later stage [26]. It is consistent with the results of this study that iSN04 treatment and subsequent suppression of Wnt/β-catenin signaling from day 5 facilitated myocardial differentiation of mESCs and miPSCs (Figure 7).

**Table 1.**
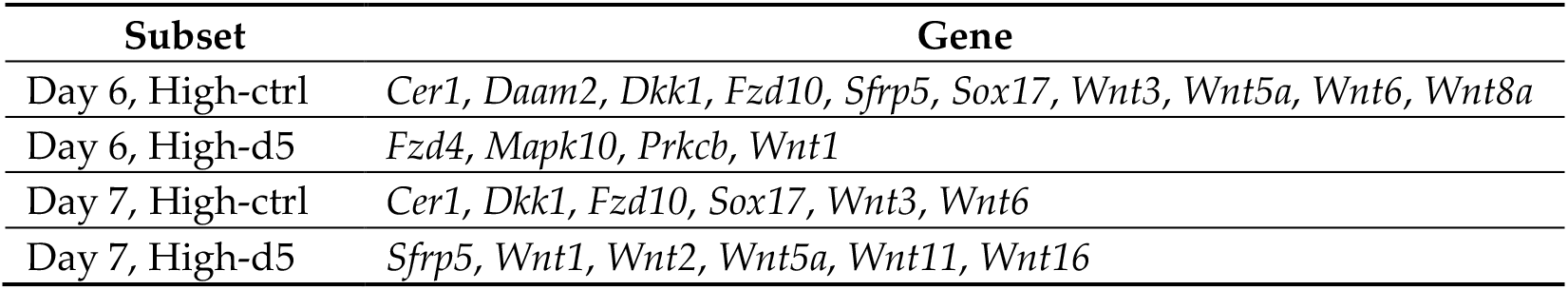
The DEGs significantly enriched in the Wnt signaling pathway.

| Subset | Gene |
| --- | --- |
| Day 6, High-ctrl | *Cer1*, *Daam2*, *Dkk1*, *Fzd10*, *Sfrp5*, *Sox17*, *Wnt3*, *Wnt5a*, *Wnt6*, *Wnt8a* |
| Day 6, High-d5 | *Fzd4*, *Mapk10*, *Prkcb*, *Wnt1* |
| Day 7, High-ctrl | *Cer1*, *Dkk1*, *Fzd10*, *Sox17*, *Wnt3*, *Wnt6* |
| Day 7, High-d5 | *Sfrp5*, *Wnt1*, *Wnt2*, *Wnt5a*, *Wnt11*, *Wnt16* |

**Figure 4.**
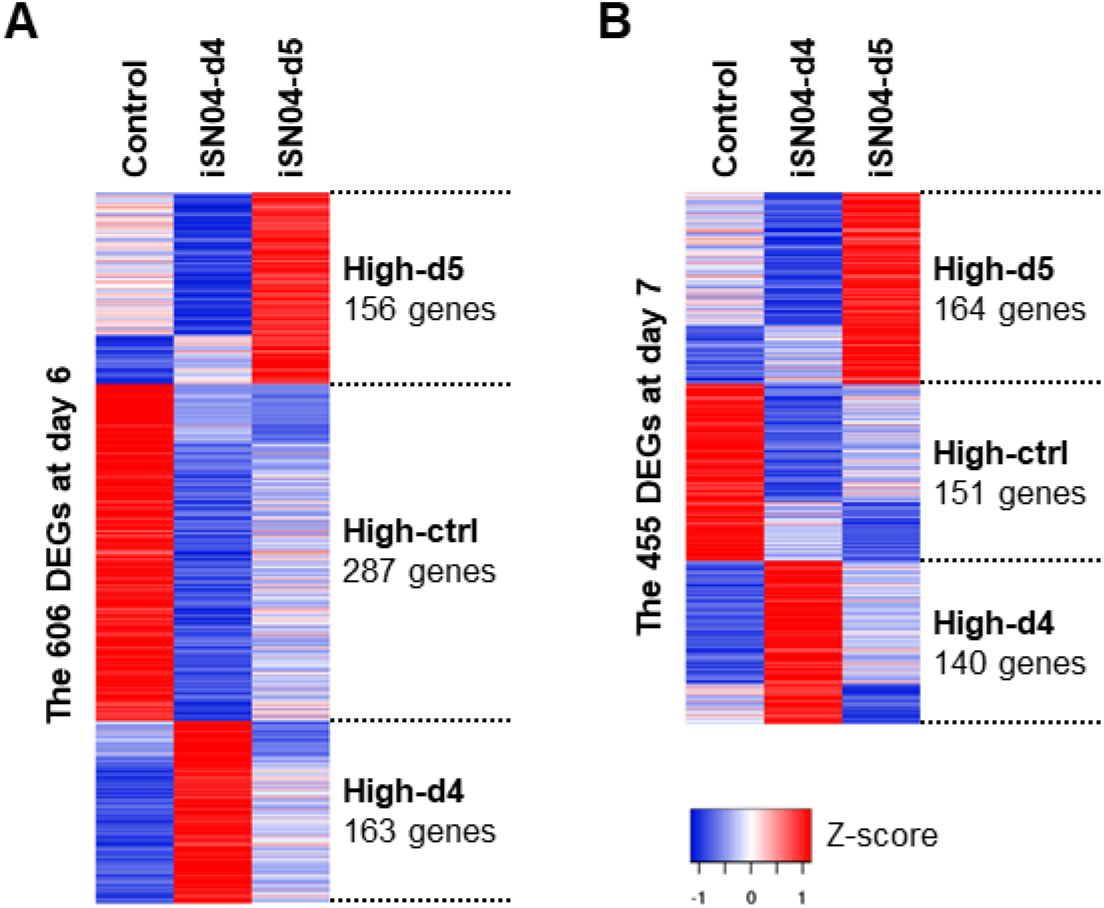
Heatmap of the iSN04-dependent DEGs in mESCs. The DEGs and their subsets at day 6 (**A**) and day 7 (**B**) in hCGp7 cells induced to differentiate in DM and treated with iSN04 from day 4 or 5 on 30-mm dishes (same samples as in Figure 2C) are shown with expression levels as Z-scores.

iSN04 also affected the levels of *Wnt2, Wnt5a*, and *Wnt11*, which are involved in the non-canonical β-catenin-independent Wnt signaling pathway. *Wnt5a* was downregulated by iSN04 (included in the High-ctrl group) on day6, but *Wnt2/5a/11* were upregulated on day 7 in the iSN04-d5 group. These Wnt signals have been reported to promote the differentiation of cardiac mesoderm into cardiomyocytes [23], corresponding to the current results that iSN04 treatment from day 5 promoted cardiomyogenesis (Figure 7).

### 2.5. iSN04 Restricts Nucleolin Translocation in PSCs

Nucleolin, the target of iSN04 [6], is a ubiquitous multifunctional phosphoprotein that nucleolin localizes to the nuclei and accumulates in the nucleoli of undifferentiated PSCs [13,14]. As shown in Figure 5, nucleolin was observed to remain localized in the nucleoli of hCGp7 cells until day 3 after differentiation. Nucleolin then began to diffuse into the cytoplasm from day 4 to at least day 7, similar to what has been reported during myogenic differentiation of skeletal muscle myoblasts [6,8]. Interestingly, iSN04 treatment from day 4 or 5 restricted nucleolin in the nucleoli and inhibited its cytoplasmic translocation, probably because iSN04 interferes with nucleolin binding to its transporter. These results suggest that nucleolar nucleolin is involved in the alteration of gene expression patterns by iSN04.

**Figure 5.**
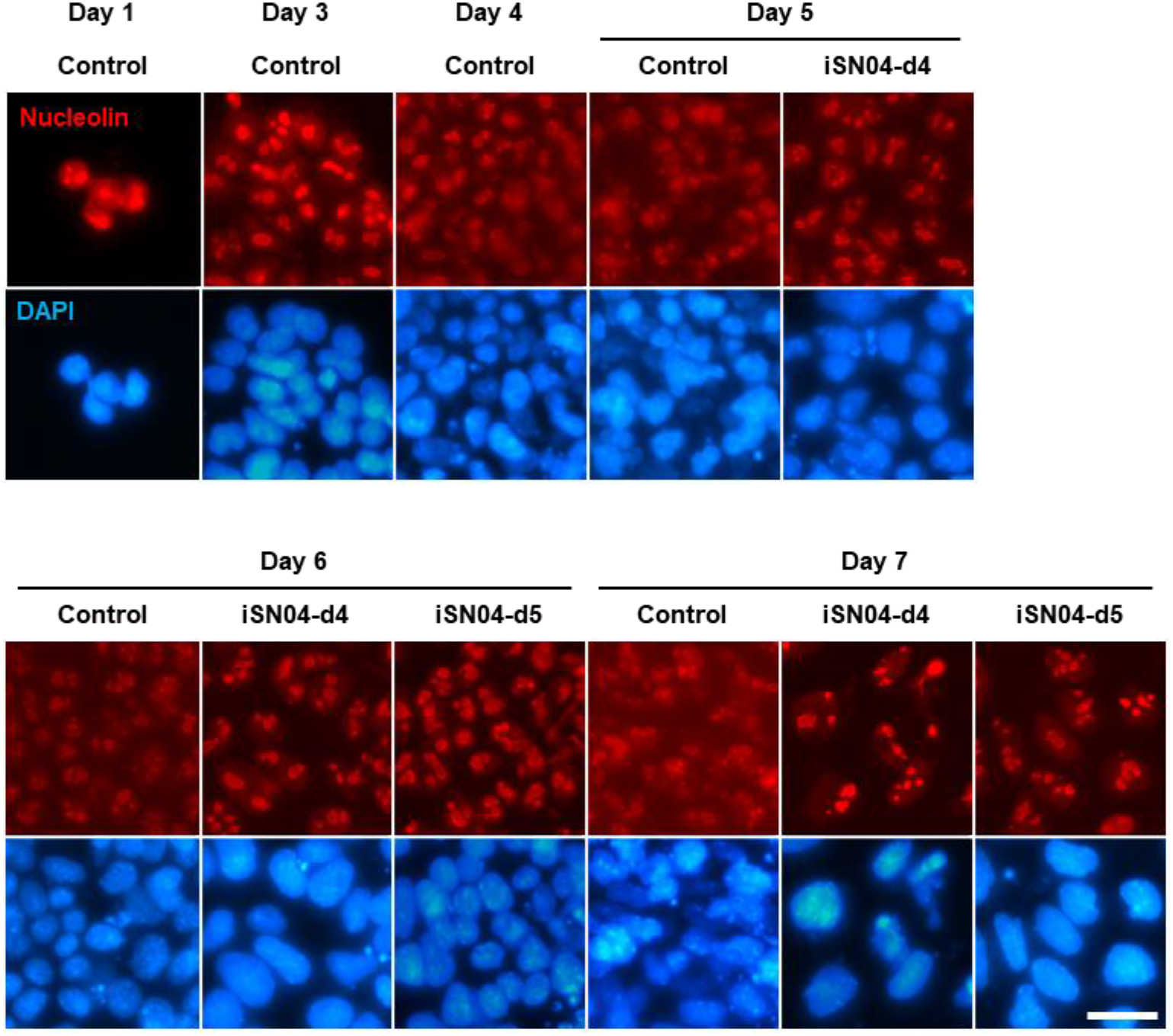
iSN04 restricts nucleolin localization in mESCs during differentiation. Representative fluorescence images of nucleolin staining of hCGp7 cells induced differentiation in DM and treated with 10 μM iSN04 from day 4 or 5 on 30-mm dishes. Scale bar, 25 μm.

### 2.6. iSN04 Does Not Affect Myocardial Cell Hypertrophy

Embryonal cardiac gene expression that arises during myocardial differentiation of PSCs is also driven in cardiomyocytes in response to various pathophysiological stimuli, leading to compensated hypertrophy and ultimately to heart failure [27,28]. Therefore, it is of concern that iSN04, which induces cardiomyogenesis, may initiate myocardial cell hypertrophy by upregulating embryonal cardiac genes. To evaluate the safety of iSN04 on cardiomyocytes, primary-cultured neonatal rat ventricular cardiomyocytes were treated with iSN04 in the presence or absence of an α_1_-adrenergic agonist, phenylephrine (PE), which induces hypertrophic responses. As shown in Figure 6, PE significantly increased the surface area of cardiomyocytes, but iSN04 had no effect on cell size regardless of PE treatment. This indicates that iSN04 does not affect the pathophysiological hypertrophy of terminally differentiated cardiomyocytes.

**Figure 6.**
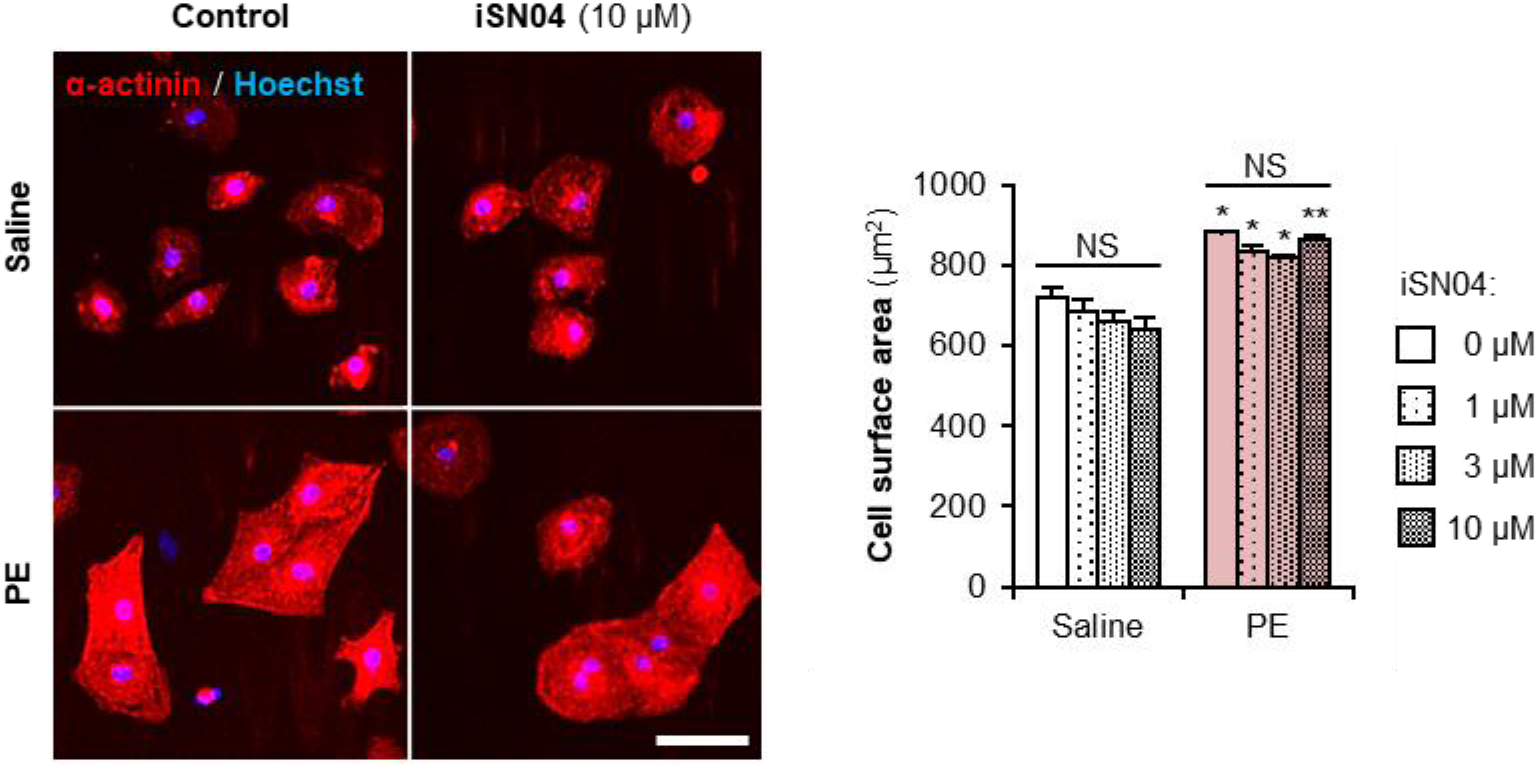
iSN04 does not affect myocardial cell hypertrophy. Representative fluorescence images of α-actinin staining of rat cardiomyocytes induced hypertrophy using 30 μM PE and treated with 10 μM iSN04 for 48 h. Scale bar, 50 μm. α-actinin^+^ cell surface area was quantified. * *p* < 0.05, ** *p* < 0.01 vs saline with the same concentration of iSN04 (Scheffe’s *F* test). NS, no significant difference. *n* =3 (200 cardiomyocytes in each experiment).

**Figure 7.**
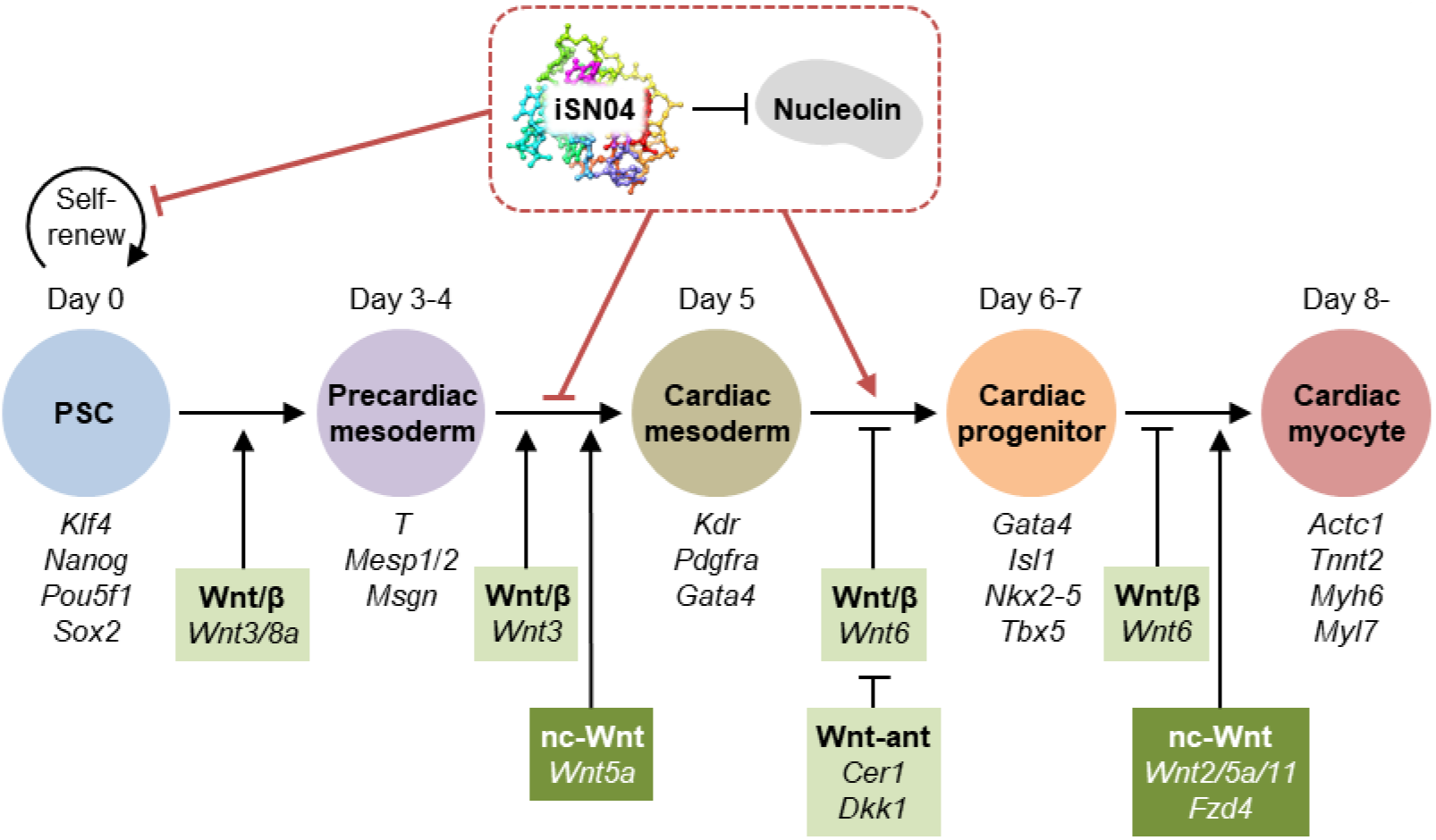
Hypothetical model of the effect of iSN04 on the Wnt signaling pathway during myocardial differentiation of PSCs. The genes investigated in this study are shown. β, β-catenin; ant, antagonist; nc, non-canonical.

## 3. Discussion

This is the first report on the generation of cardiomyocytes from PSCs using a DNA aptamer. iSN04 was originally identified as a myogenetic oligodeoxynucleotide that induces myogenic differentiation of skeletal muscle myoblasts [6-9]. iSN04 serves as an anti-nucleolin aptamer that inhibits proliferation and induces differentiation not only of myoblasts but also of rhabdomyosarcoma, a soft tissue tumor of the striated muscle [10]. Since downregulation of nucleolin has been reported to impair the growth of mESCs [13,15,16], we hypothesized that iSN04 would inhibit the proliferation of PSCs and enhance the production of skeletal muscle cells. As expected, the present study showed that iSN04 suppressed the growth of murine PSC colonies in the undifferentiated state. However, in the differentiating condition, iSN04 treatment from day 5 onward dramatically generated beating mature cardiomyocytes, whereas treatment from earlier stages completely inhibited cardiomyogenesis. The effect of iSN04 on skeletal muscle differentiation of PSCs was not confirmed in this study. For example, *Myod1* (MyoD) and *Myog* (myogenin), specific myogenic regulatory factors essential for skeletal myogenesis, were not detected by RNA-seq (FPKM < 1.0) in any sample throughout the differentiation period. This may be because skeletal muscle formation occurs after heart development. During murine embryogenesis, cardiac mesoderm begins to migrate at embryonic day 7 (E7) to form the heart tube at E8.5, and the major structures of the heart are established at E13.5 [29]; whereas skeletal muscle development begins at E8.5/9 and continues until birth (E19) [30]. In murine PSC culture, terminally differentiated cardiomyocytes can be generated with relatively easily in about one week as shown in this study, but skeletal muscle stem cells are difficult to obtain and require 2-3 weeks even with the optimized protocol [31]. Therefore, the effect of iSN04 on PSCs might first appear in cardiomyogenesis. The myogenic capacity of iSN04 during skeletal muscle differentiation of PSCs needs to be investigated in further studies under more specific culture conditions and treatment duration.

Gene expression analysis by qPCR and RNA-seq revealed that nucleolin inhibition by iSN04 in mESCs from day 3-4 completely inhibited the differentiation into cardiac mesoderm, but that from day 5 significantly improved the myocardial differentiation into cardiomyocytes. The iSN04-dependent DEGs suggested that the Wnt signaling pathway is involved in this stage-specific effect of iSN04. The role of Wnt signaling during cardiomyogenesis of PSCs is quite complicated. As shown in Figure 7, Wnt/β-catenin signaling directs cardiac commitment in the early differentiation stages, but subsequently blocks terminal differentiation into cardiomyocytes [21,23,24,32]. In contrast, non-canonical Wnt signaling consistently promotes myocardial differentiation from precardiac mesoderm [23]. The RNA-seq data indicated that iSN04 suppressed the Wnt/β-catenin signaling pathway and upregulated non-canonical Wnt signals as a mechanism of the time-dependent effect of iSN04. Nucleolin is known to promote the Wnt/β-catenin signaling. Normally, free β-catenin is phosphorylated by glycogen synthase kinase 3β (GSK-3β) and then degraded by the proteasome. Upon various stimuli, nucleolin enhances GSK-3β phosphorylation to inactivate it, resulting in an increase in β-catenin protein [33]. Therefore, antagonizing nucleolin with iSN04 reduces β-catenin protein levels and then inhibits its nuclear translocation and downstream transcription [34]. On the other hand, the relationship between nucleolin and the non-canonical Wnt signaling pathway is still unclear. Immunostaining visualized that iSN04 treatment retained nucleolin in nucleoli during the differentiation of PSCs. Since nucleolar nucleolin is involved in chromatin remodeling, gene transcription, and RNA metabolism [11], iSN04-anchored nucleolin may contribute to the alteration of gene expression patterns, including non-canonical Wnt signaling.

PSC-derived cardiomyocytes are expected to be cell sources for regenerative therapy of heart failure, which is the final stage of cardiac diseases. Cardiomyocytes in the overloaded heart undergo hypertrophy and eventually become dysfunctional. During this process, the gene expression profile in the myocardium shifts from the adult state to the embryonic type, including at the epigenetic level [27,28]. There was a potential risk that iSN04 would induce hypertrophic responses of cardiomyocytes in addition to cardiomyogenesis of PSCs. In practice, however, iSN04 did not affect PE-induced pathophysiological hypertrophy of primary-cultured cardiomyocytes. A recent study reported that inhibition of the Wnt/β-catenin/GSK-3β pathway ameliorates myocardial cell hypertrophy [35]. If iSN04 also suppresses Wnt/β-catenin signaling in cardiomyocytes, iSN04 would not have the risk of inducing hypertrophic responses. In cardiac regeneration therapy, PSC-derived cardiac progenitors or immature cardiomyocytes are transplanted into the heart and induced to regenerate functional myocardium [21]. iSN04 may be a potentially safe and useful molecule for the generation of PSC-derived cardiomyocytes in vitro and in vivo for cardiac tissue reconstruction.

This study demonstrated that iSN04 induced myocardial differentiation of two murine PSC lines, hCGp7 mESCs and 20D17 miPSCs. The amino acid sequence of nucleolin is highly conserved between mouse and human (83.0% identity and 89.0% similarity). Since iSN04 has indeed enhanced the skeletal muscle differentiation of both murine and human myoblasts [6,7,9], it is promising that iSN04 will interfere with nucleolin in human PSCs. Murine and human PSCs differ in their requirement for leukemia inhibitory factor (LIF), in the morphology of the undifferentiated cell colony, and in the culture method for cardiomyogenesis [32]. In order to apply iSN04 for regenerative therapy in clinical settings, its effect on human PSCs needs to be validated by further studies. The establishment of iSN04-induced cardiomyocytes from PSCs will provide the alternative technology for heart regeneration in the future.

## 4. Materials and Methods

### 4.1. Chemicals

iSN04 (5’-AGA TTA GGG TGA GGG TGA-3’), in which all phosphodiester bonds were phosphorothioated to enhance nuclease resistance, was synthesized and HPLC-purified (GeneDesign, Osaka, Japan) and then dissolved in endotoxin-free water [6]. PE (Fujifilm Wako Chemicals, Osaka, Japan) was dissolved in saline. Equal volumes of the solvents were used as the negative controls.

### 4.2. mESCs amd miPSCs

The experimental procedure for the preparation of MEFs was performed in accordance with the Regulations for Animal Experimentation of Shinshu University, and the animal protocol was approved by the Committee for Animal Experiments of Shinshu University (No. 280083). MEFs were prepared from E12 embryos of Slc:ICR mice (Japan SLC, Shizuoka, Japan) [6]. The embryos, from which the heads and internal organs were removed, were minced in DMEM (Nacalai, Osaka, Japan) containing 10% fetal bovine serum (FBS) (HyClone; Cytiva, Marlborough, MA, USA) and a mixture of 100 units/ml penicillin and 100 μg/ml of streptomycin (P/S) (Nacalai). The tissue clusters were cultured for 3 days, then the outgrowing cells were dissociated into single cells as MEFs by 0.25% trypsin with 1 mM EDTA (Fujifilm Wako Chemicals). The MEFs seeded on fresh dishes were treated with 10 μg/ml MMC (Fujifilm Wako Chemicals) for 2 h, then the cells were frozen and stored at -80°C until use. The MMC-MEFs were seeded on gelatin-coated dishes as a feeder layer for mESCs and miPSCs.

The miPSC line, 20D17, in which a GFP-IRES-Puro^r^ cassette was inserted into the 5’ UTR of the *Nanog* gene [17], was provided by the RIKEN BRC (Tsukuba, Japan) through the Project for Realization of Regenerative Medicine and the National Bio-Resource Project of the MEXT, Japan. The 129/Ola-derived mESC line, hCGp7, in which an EGFP-PGKPuro^r^ cassette was inserted into the *Nkx2-5* locus [18,19]. Undifferentiated miPSCs and mESCs were maintained on a feeder layer in GM consisting of DMEM, 15% FBS, 1% non-essential amino acids (NEAA) (Fujifilm Wako Chemicals), 1% nucleosides (Merck, Darmstadt, Germany), 0.1 mM 2-mercaptoethanol (Nacalai), 10^3^ units/ml LIF (Nacalai), 3 μM CHIR99021 (Fujifilm Wako Chemicals), 0.8 μM PD0325901 (Fujifilm Wako Chemicals), and P/S [16]. For colony formation, fully dissociated undifferentiated 20D17 or hCGp7 cells were seeded on a feeder layer in GM. The next day, the medium was changed to fresh GM containing 10 μM iSN04. After 24 or 48 h, the cell colonies were subjected to fluorescent imaging, ALP staining, or RNA extraction. For spontaneous differentiation, undifferentiated 20D17 cells or hCGp7 cells were seeded on feeder-free gelatin-coated 30-mm dishes (3.0 × 10^4^ cells/dish) or 96-well plates (2.0 × 10^3^ cells/well) in DM consisting of DMEM, 10% FBS, 5% horse serum (HyClone; Cytiva), 1% NEAA, 0.1 mM 2-mercaptoethanol, and P/S (defined as day 0) [31,36]. DM (containing 10 μM iSN04 if necessary) was changed every two days. All cells were cultured at 37°C under 5% CO_2_ throughout the experiments.

### 4.3. Cell Imaging

Fluorescence and phase-contrast images were captured using EVOS FL Auto microscope (AMAFD1000; Thermo Fisher Scientific, Waltham, MA, USA). Colony size of mESCs was measured using ImageJ software version 1.52a (Wayne Rasband; National Institute of Health, Bethesda, MD, USA). Phase-contrast videos were captured using CKX53 microscope (Olympus, Tokyo, Japan) with Moticam 1080 digital camera system (Shimadzu RIKA Corporation, Tokyo, Japan).

### 4.4. ALP Staining

ALP enzymatic activity of the hCGp7 cell colonies was visualized using ALP Stain Kit (Fujifilm Wako Chemicals) according to the manufacturer’s instructions [37].

### 4.5. Nucleolin Staining

hCGp7 cells were fixed with 2% paraformaldehyde, permeabilized with 0.2% Triton X-100, and immunostained with 1.0 μg/ml rabbit polyclonal anti-nucleolin antibody (ab22758; Abcam, Cambridge, UK) overnight at 4°C and then with 0.1 μg/ml Alexa Fluor 594-conjugated donkey polyclonal anti-rabbit IgG antibody (Jackson ImmunoResearch, West Grove, PA, USA) for 1 h at room temperature [6-8,10]. Cell nuclei were stained with DAPI (Nacalai).

### 4.6. qPCR

Total RNA from hCGp7 cells was isolated using NucleoSpin RNA Plus (Macherey-Nagel, Düren, Germany) and was reverse transcribed using ReverTra Ace qPCR RT Master Mix (TOYOBO, Osaka, Japan). qPCR was performed using GoTaq qPCR Master Mix (Promega, Madison, WI, USA) with StepOne Real-Time PCR System (Thermo Fisher Scientific). The amount of each transcript was normalized to that of tyrosine 3-monooxygenase/tryptophan 5-monooxygenase activation protein zeta (*Ywhaz*) gene. Results are presented as fold-change. Primer sequences are listed in Supplementary Table S1.

### 4.7. RNA-Seq

Total RNA from hCGp7 cells used for qPCR was subjected to RNA-seq (Novogene, Beijing, China). RNA qualities were checked using Agilent 2100 Bioanalyzer (Agilent Technologies, Waldbronn, Germany). RNA integrity number (RIN) values were 10.0 in all samples. The RNA was subjected to prepare libraries using Illumina TruSeq RNA and DNA Sample Prep Kits (Illumina, San Diego, CA, USA). Library qualities were confirmed using Qubit 2.0 fluorometer (Life Technologies; Thermo Fisher Scientific) and Agilent 2100 Bioanalyzer. RNA-seq was performed using Illumina NovaSeq 6000 (Illumina) to generate > 6-GB raw data per sample. Raw data was recorded in a FASTQ format. The quality of the read was calculated as the arithmetic mean of its Phred quality scores. Then the reads as follows were discarded; with adaptor contamination, when uncertain nucleotides constitute > 10% of either read, or when low quality nucleotides (base quality < 20) constitute > 50% of the read. The cleaned reads were used for subsequent analyses. The reads were mapped to a reference genome GRCm39 using TopHat2. The numbers of the reads and mapping efficiencies were summarized in Supplementary Table S2. The expression levels of the transcripts were calculated as FPKM using HTSeq.

### 4.8. PCA

The FPKM values (> 1.0) of the RNA-seq data were subjected to PCA. A total of 13,613 genes were used as the number of dimensions for the 9 sample vectors. Each vector contained the FPKM values as the elements. Variance-covariance matrices (13,613 × 13,613 dimension) were calculated from the 9 vectors. The matrices were diagonalized, and the eigenvalues and eigenvectors were obtained. The projection onto the eigenvector of the largest eigenvalue corresponded to the first, second, and third components of the PCA (PC1, PC2, and PC3). These calculations were performed using Python scripts [38].

### 4.9. Heatmap and KEGG Pathway Analysis

Heatmap of the expression levels of the DEGs was generated by Heatmapper (http://www.heatmapper.ca/) [39] with following settings. Clustering method, average linkage; distance measurement method, Spearman rank correlation. Each row represents one gene, and each column represents one sample. The Z-score representing the red and blue gradients indicates an increase and a decrease in gene expression, respectively. The DEGs were subjected to KEGG pathway analysis using DAVID Bioinformatics Resources 6.8 (https://david.ncifcrf.gov/) [40]. The KEGG pathway with *p* value < 0.05 was defined as the signaling pathway containing significantly enriched genes.

### 4.10. Cardiomyocytes

The experimental procedure for the preparation of rat cardiomyocytes was conducted in accordance with the Guide for the Care and Use of Laboratory Animals published by the University of Shizuoka, and the animal protocol was approved by the Ethics Committee of the University of Shizuoka (No. US176278). Neonatal rat ventricular cardiomyocytes were isolated from 1-day-old Sprague-Dawley rats (Japan SLC) and seeded on 24-well plates (5.0 × 10^4^ cells/well) as previously described [41-43]. After 36 h, the cardiomyocytes were treated with 1-10 μM iSN04 for 2 h and then hypertrophic responses were induced by 30 μM PE for 48 h with continued iSN04 treatment. The cells were subjected to immunocytochemistry using mouse anti-α-actinin antibody (Sigma-Aldrich, Saint Louis, MO, USA), Alexa Fluor 555-conjugated goat anti-mouse IgG antibody (Thermo Fisher Scientific), and Hoechst 33258 (Dojindo, Kumamoto, Japan). Fluorescent images were captured and cell surface area of 200 α-actinin^+^ cardiomyocytes per sample was automatically measured using ArrayScan (Thermo Fisher Scientific) [43].

### 4.11. Statistical Analysis

Results are presented as mean ± standard error. Statistical comparisons between two groups were performed using unpaired two-tailed Student’s *t*-test and among multiple groups using Tukey-Kramer test or Scheffe’s *F* test after one-way analysis of variance, as appropriate. Statistical significance was set at *p* < 0.05. Correlation analysis was performed using Pearson’s correlation coefficient test.

## 5. Conclusions

An anti-nucleolin aptamer, iSN04, suppressed the proliferation of undifferentiated murine PSCs. In differentiating conditions, iSN04 treatment at earlier stages completely inhibited to lead cardiac lineage, but that at later stages significantly enhanced the myocardial differentiation into cardiomyocytes. This stage-specific effect of iSN04 during cardiomyogenesis of PSCs is, in part, due to modulation of the Wnt signaling pathway. Since iSN04 did not affect hypertrophic responses in cardiomyocytes, iSN04 may be a useful molecule to generate PSC-derived cardiomyocytes for heart regeneration therapy.

## 6. Patents

T.T. is the inventor of Japanese Patent No. 7152001 covering iSN04-induced myocardial differentiation of PSCs.

## Supplementary Materials

Figure S1: Correlation of the gene expression levels quantified by RNA-seq and qPCR, Figure S2: PCA of the RNA-seq data, Table S1: Primer sequences for qPCR, Table S2: The numbers of the reads obtained by RNA-seq, Video S1: Beating cardiomyocytes spontaneously differentiated from miPSCs, Video S2: Beating cardiomyocytes differentiated from miPSCs treated with iSN04. References [44-46] are cited in the Supplementary Materials.

## Author Contributions

T.T. designed the study and drafted the manuscript. M.I., Y.N., Y.S., and K.U. performed the experiments and data analyses. T.S., H.K., and T.M. provided the materials. All authors have read and approved to the published version of the manuscript.

## Funding

This research was funded by the Japan Society for the Promotion of Science (19K05948 and 22K05554) to T.T. and by the Fund of Nagano Prefecture to Promote Scientific Activity (R1-3-3) to M.I.

## Informed Consent Statement

Not applicable.

## Data Availability Statement

FASTQ raw read data of the RNA-seq is deposited in DDBJ Sequence Read Archive (Research Organization of Information and Systems, National Institute of Genetics, Mishima, Japan) with the accession number: DRA016771. Other raw data supporting the conclusions of this article will be made available by the authors, without undue reservation.

## Conflicts of Interest

The authors declare no conflict of interest.

## Abbreviations

ALP: Alkaline phosphatase
DEG: Differentially expressed gene
DM: Differentiation medium
ESC: Embryonic stem cell
FPKM: Fragments per kilobase per million reads
GFP: Green fluorescent protein
GM: Growth medium
iPSC: Induced pluripotent stem cell
MEF: Murine embryonic fibroblast
MMC: Mitomycin C
myoDN: Myogenetic oligodeoxynucleotide
PCA: Principal component analysis
PE: Phenylephrine
PSC: Pluripotent stem cell
qPCR: Quantitative real-time RT-PCR
RNA-seq: RNA sequencing

## Supplementary Materials

**Supplementary Figure S1.**
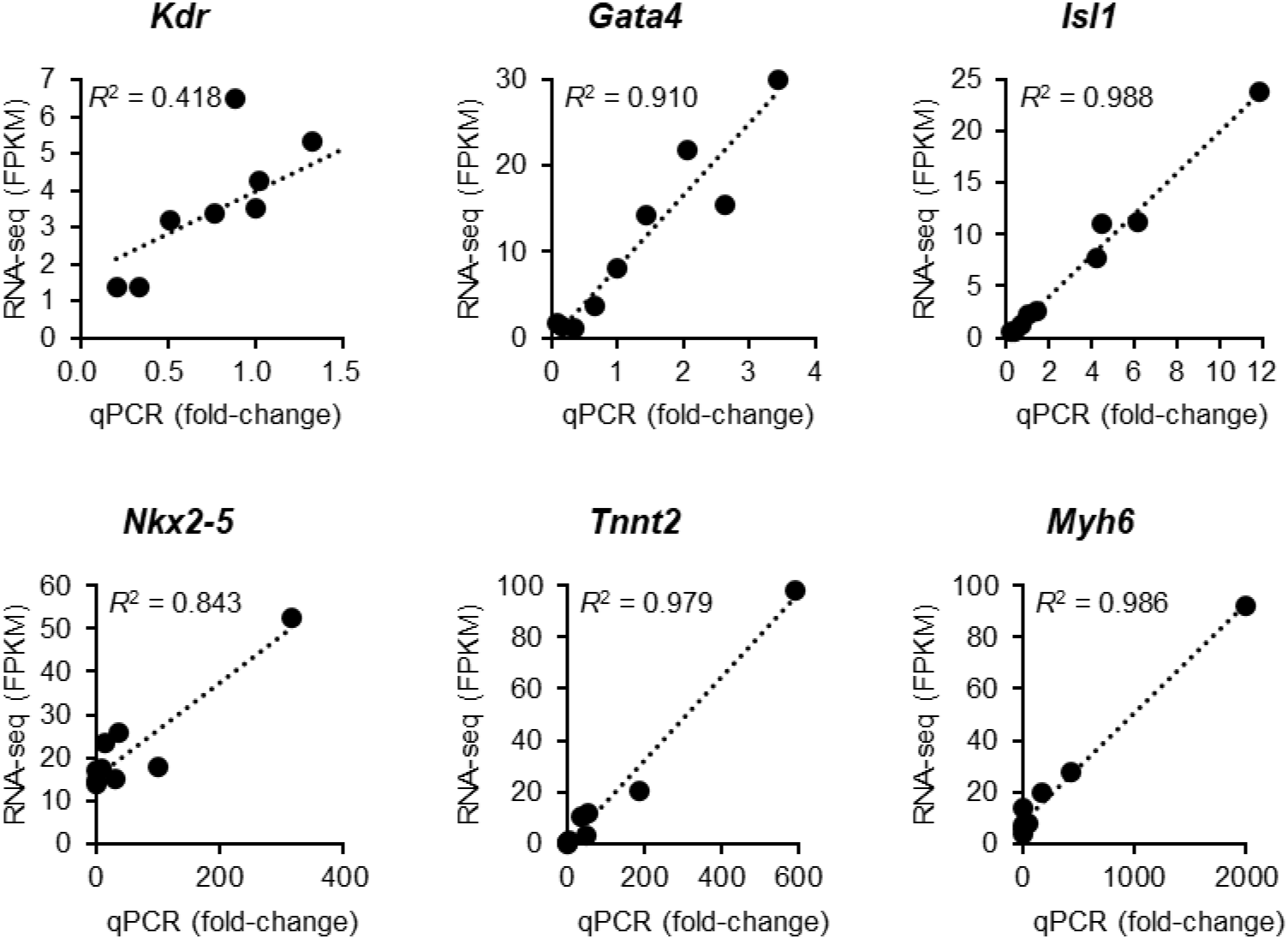
Correlation of the gene expression levels quantified by RNA-seq and qPCR. The FPKM values defined by RNA-seq and the mRNA levels (fold-changes) detected by qPCR (Figure 2C) were analyzed using Pearson’s correlation coefficient test.

**Supplementary Figure S2.**
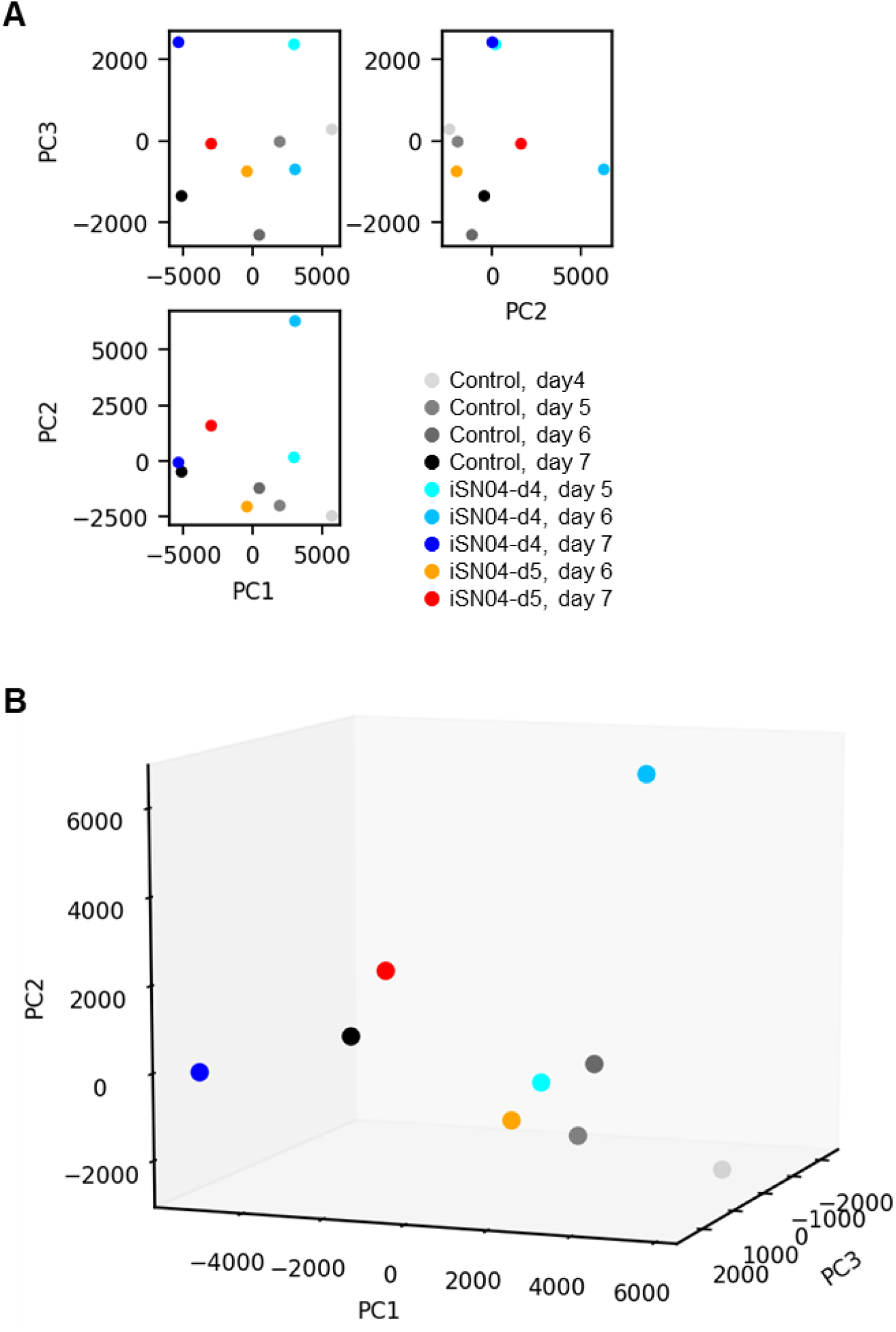
PCA of the RNA-seq data. The 13,613 FPKM values obtained by RNA-seq of hCGp7 cells induced to differentiate in DM and treated with 10 μM iSN04 from day 4 or 5 on 30-mm dishes (same samples as in Figure 2C). (**A**) Two-dimensional plots on the PCA space reconstructed by PC1 (contribution, 0.52), PC2 (0.26), and PC3 (0.09). (**B**) Three-dimensional plot on the PCA space reconstructed by PC1, PC2, and PC3.

**Supplementary Table S1.**
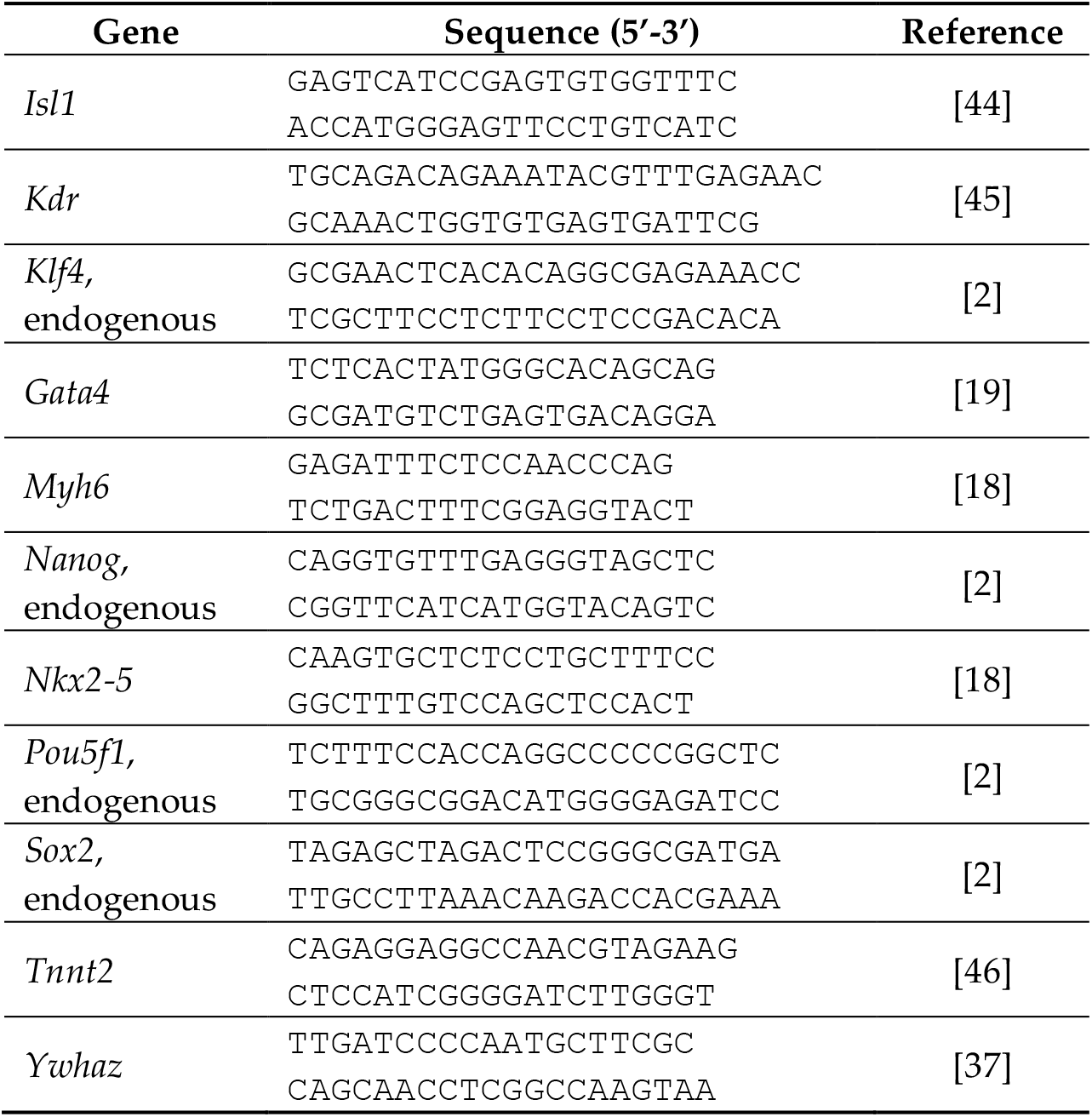
Primer sequences for qPCR.

**Supplementary Table S2.** The numbers of the reads obtained by RNA-seq.

| Sample | Raw reads | Clean reads | Mapped reads | % mapped |
| --- | --- | --- | --- | --- |
| Control, day 4 | 50,439,826 | 497,66,540 | 46,032,264 | 92.50% |
| Control, day 5 | 54,721,348 | 54,064,028 | 50,082,857 | 92.64% |
| Control, day 6 | 54,782,002 | 54,174,286 | 50,116,711 | 92.51% |
| Control, day 7 | 48,692,658 | 48,064,434 | 44,481,973 | 92.55% |
| iSN04-d4, day 5 | 59,925,566 | 59,225,592 | 54,811,573 | 92.55% |
| iSN04-d4, day 6 | 52,889,966 | 52,342,122 | 47,145,750 | 90.07% |
| iSN04-d4, day 7 | 60,513,932 | 59,768,940 | 55,336,829 | 92.58% |
| iSN04-d5, day 6 | 56,021,784 | 55,366,368 | 51,361,265 | 92.77% |
| iSN04-d5, day 7 | 55,013,818 | 54,174,286 | 50,213,955 | 92.51% |

**Supplementary Video S1**. Beating cardiomyocytes spontaneously differentiated from miPSCs. Un-differentiated 20D17 cells were seeded on gelatin-coated dishes and induced spontaneous differentiation in DM for 11 days.

**Supplementary Video S2**. Beating cardiomyocytes differentiated from miPSCs treated with iSN04. Undifferentiated 20D17 cells were seeded on gelatin-coated dishes and induced spontaneous differentiation in DM for 11 days. The cells were treated with 10 μM iSN04 from day 5 to day 9.

## Notes

### Competing Interest Statement

The authors have declared no competing interest.

## References

1. Dogan, A. Embryonic stem cells in development and regenerative medicine. Adv. Exp. Med. Biol. 2018, 1079, 1–15.

2. Takahashi, K.; Yamanaka, S. Induction of pluripotent stem cells from mouse embryonic and adult fibroblast cultures by defined factors. Cell 2006, 126, 663–676.

3. Takahashi, K.; Tanabe, K.; Ohnuki, M.; Narita, M.; Ichisaka, T.; Tomoda, K.; Yamanaka, S. Induction of pluripotent stem cells from adult human fibroblasts by defined factors. Cell 2007, 131, 861–872.

4. Wang, T.; Chen, C.; Larcher, L.M.; Barrero, R.A.; Veedu, R.N. Three decades of nucleic acid aptamer technologies: Lessons learned, progress and opportunities on aptamer development. Biotechnol. Adv. 2019, 37, 28–50.

5. Ueki, R.; Atsuta, S.; Ueki, A.; Hoshiyama, J.; Li, J.; Hayashi, Y.; Sando, S. DNA aptamer assemblies as fibroblast growth factor mimics and their application in stem cell culture. Chem. Commun. 2019, 55, 2672–2675.

6. Shinji, S.; Umezawa, K.; Nihashi, Y.; Nakamura, S.; Shimosato, T.; Takaya, T. Identification of the myogenetic oligodeoxynucleotides (myoDNs) that promote differentiation of skeletal muscle myoblasts by targeting nucleolin. Front. Cell Dev. Biol. 2021, 8, 616706.

7. Nakamura, S.; Yonekura, S.; Shimosato, T.; Takaya, T. Myogenetic oligodeoxynucleotide (myoDN) recovers the differentiation of skeletal muscle myoblasts deteriorated by diabetes mellitus. Front. Physiol. 2021, 12, 679152.

8. Nihashi, Y.; Shinji, S.; Umezawa, K.; Shimosato, T.; Ono, T.; Kagami, H.; Takaya, T. Myogenetic oligodeoxynucleotide complexed with berberine promotes differentiation of chicken myoblasts. Anim. Sci. J. 2021, 92, e13597.

9. Nihashi, Y.; Yamamoto, M.; Shimosato, T.; Takaya, T. Myogenetic oligodeoxynucleotide restores differentiation and reverses inflammation of myoblasts aggravated by cancer-conditioned medium. Muscles 2022, 1, 111–120.

10. Nohira, S.; Shinji, S.; Nakamura, Y.; Nihashi, Y.; Shimosato, T.; Takaya, T. Myogenetic oligodeoxynucleotides as anti-nucleolin aptamers inhibit the growth of embryonal rhabdomyosarcoma cells. Biomedicines 2022, 10, 2691.

11. Jia, W.; Yao, Z.; Zhao, J.; Guan, Q.; Gao, L. New perspectives of physiological and pathological functions of nucleolin (NCL). Life Sci. 2017, 186, 1–10.

12. Laurincik, J.; Bjerregaard, B.; Strejcek, F.; Rath, D.; Niemann, H.; Rosenkranz, C.; Ochs, R.L.; Maddox-Hyttel, P. Nucleolar ultrastructure and protein allocation in in vitro produced porcine embryos. Mol. Reprod. Dev. 2004, 68, 327–334.

13. Li, H.; Wang, B.; Yang, A., Lu, R.; Wang, W.; Zhou, Y.; Shi, G.; Kwon, S.W.; Zhao, Y.; Jin, Y. Ly-1 antibody reactive clone is an important nucleolar protein for control of self-renewal and differentiation in embryonic stem cells. Stem Cells 2009, 27, 1244–1254.

14. Johansson, H.; Svensson, F.; Runnberg, R.; Simonsson, T.; Simonsson, S. Phosphorylated nucleolin interacts with translationally controlled tumor protein during mitosis and with Oct4 during interphase in ES cells. PLoS One 2010, 5, e13678.

15. Yang, A.; Shi, G.; Zhou, C.; Lu, R.; Li, H.; Sun, L.; Jin, Y. Nucleolin maintains embryonic stem cell self-renewal by suppression of p53 protein-dependent pathway. J. Biol. Chem. 2011, 286, 43370–43382.

16. Percharde, M.; Lin, C.J.; Yin, Y.; Guan, J.; Peixoto, G.A.; Bulut-Karslioglu, A.; Biechele, S.; Huang, B.; Shen, X.; Ramalho-Santos, M. A LINE1-nucleolin partnership regulates early development and ESC identity. Cell 2018, 174, 391–405.

17. Okita, K.; Ichisaka, T.; Yamanaka, S. Generation of germline-competent induced pluripotent stem cells. Nature 2007, 488, 313–317.

18. Hidaka, K.; Lee, J.K.; Kim, H.S; Ihm, C.H.; Iio, A.; Ogawa, M.; Nishikawa, S.; Kodama, I.; Morisaki, T. Chamber-specific differentiation of Nkx2.5-positive cardiac precursor cells from murine embryonic stem cells. FASEB J. 2003, 17, 740–742.

19. Kaichi, S.; Takaya, T.; Morimoto, T.; Sunagawa, Y.; Kawamura, T.; Ono, K.; Shimatsu, A.; Baba, S.; Heike, T.; Nakahata, T.; Hasegawa, K. Cyclin-dependent kinase 9 forms a complex with GATA4 and is involved in the differentiation of mouse ES cells into cardiomyocytes. J. Cell. Physiol. 2011, 226, 248–254.

20. Narita, S.; Unno, K.; Kato, K.; Okuno, Y.; Sato, Y.; Tsumura, Y.; Fujikawa, Y.; Shimizu, Y.; Hayashida, R.; Kondo, K.; Shibata, R.; Murohara, T. Direct reprogramming of adult adipose-derived regenerative cells toward cardiomyocytes using six transcriptional factors. iScience 2022, 25, 104651.

21. Mercola, M.; Ruiz-Lozano, P.; Schneider, M.D. Cardiac muscle regeneration: lessons from development. Genes Dev. 2011, 25, 299–309.

22. Nusse, R.; Clevers, H. Wnt/β-catenin signaling, disease, and emerging therapeutic modalities. Cell 2017, 169, 989–999.

23. Mazzotta, S.; Neves, C.; Bonner, R.J.; Bernardo, A.S.; Docherty, K.; Hoppler, S. Distinctive roles of canonical and noncanonical Wnt signaling in human embryonic cardiomyocyte development. Stem Cell Reports 2016, 7, 764–776.

24. Schmeckpeper, J.; Verma, A.; Yin, L.; Beigi, F.; Zhang, L.; Payne, A.; Zhang, Z.; Pratt, R.E.; Dzau, V.J.; Mirotsou, M. Inhibition of Wnt6 by Sfrp2 regulates adult cardiac progenitor cell differentiation by differential modulation of Wnt pathways. J. Mol. Cell. Cardiol. 2015, 85, 215–225.

25. Mukherjee, S.; Luedeke, D.M.; McCoy, L.; Iwafuchi, M.; Zorn, A.M. SOX transcription factors direct TCF-independent WNT/beta-catenin responsive transcription to govern cell fate in human pluripotent stem cells. Cell Rep. 2022, 40, 111247.

26. Naito, T.A.; Shiojima, I.; Akazawa, H.; Hidaka, K.; Morisaki, T.; Kikuchi, A., Komuro, I. Developmental stage-specific biphasic roles of Wnt/β-catenin signaling in cardiomyogenesis and hematopoiesis. Proc. Natl. Acad. Sci. USA 2006, 103, 19812–19817.

27. Frey, N.; Katus, H.A.; Olson, E.N.; Hill, J.A. Hypertrophy of the heart: a new therapeutic target? Circulation 2004, 109, 1580–1589.

28. Lei, H.; Hu, J.; Sun, K.; Xu, D. The role and molecular mechanism of epigenetics in cardiac hypertrophy. Heart Fail. Rev. 2020, doi: 10.1007/s10741-020-09959-3.

29. D’Amato, G.; Luxan, G.; de la Pmpa, J.L. Notch signalling in ventricular chamber development and cardiomyopathy. FEBS J. 2016, 283, 4223–4237.

30. Tajbakhsh, S. Skeletal muscle stem cells in developmental versus regenerative myogenesis. J. Intern. Med. 2009, 266, 372–389.

31. Mizuno, Y.; Chang, H.; Umeda, K.; Niwa, A.; Iwasa, T.; Awaya, T.; Fukada, S.; Yamamoto, H.; Yamanaka, S.; Nakahata, T.; Heike, T. Generation of skeletal muscle stem/progenitor cells from murine induced pluripotent stem cells. FASEB J. 2010, 24, 2245–2253.

32. Mummery, C.L.; Zhang, J.; Ng, E.S.; Elliott, D.A.; Elefanty, A.G.; Kamp, T.J. Differentiation of human embryonic stem cells and induced pluripotent stem cells to cardiomyocytes: a methods overview. Circ. Res. 2012, 111, 344–358.

33. Reister, S.; Mahotka, C.; van den Hofel, N.; Grinstein, E. Nucleolin promotes Wnt signaling in human hematopoietic stem/progenitor cells. Leukemia 2019, 33, 1052–1054.

34. Yamamoto, M.; Miyoshi, M.; Morioka, K.; Mitani, T.; Takaya, T. Anti-nucleolin aptamer, iSN04, inhibits the inflammatory responses in C2C12 myoblasts by modulating the β-catenin/NF-κB signaling pathway. Biochem. Biophys. Res. Commun. 2023, 664, 1–8.

35. Liu, J.J.; Shentu, L.M.; Ma, N.; Wang, L.Y.; Zhang, G.M.; Sun, Y.; Wang, Y.; Li, J.; Mu, Y.L. Inhibition of NF-κB and Wnt/β-catenin/GSK3β signaling pathways ameliorates cardiomyocyte hypertrophy and fibrosis in streptozotocin (STZ)-induced type 1 diabetic rats. Curr. Med. Sci. 2020, 40, 35–47.

36. Takaya, T.; Ono, K.; Kawamura, T.; Takanabe, R.; Kaichi, S.; Morimoto, T.; Wada, H.; Kita, T.; Shimatsu, A.; Hasegawa, K. MicroRNA-1 and microRNA-133 in spontaneous myocardial differentiation of mouse embryonic stem cells. Circ. J. 2009, 73, 1492–1497.

37. Nihashi, Y.; Miyoshi, M.; Umezawa, K.; Shimosato, T.; Takaya, T. Identification of a novel osteogenetic oligodeoxynucleotide (osteoDN) that promotes osteoblast differentiation in a TLR9-independent manner. Nanomaterials 2022, 12, 1680.

38. Nihashi, Y.; Umezawa, K.; Shinji, S.; Hamaguchi, Y.; Kobayashi, H.; Kono, T.; Ono, T.; Kagami, H.; Takaya, T. Distinct cell proliferation, myogenic differentiation, and gene expression in skeletal muscle myoblasts of layer and broiler chickens. Sci. Rep. 2019, 9, 16527.

39. Babicki, S.; Arndt, D.; Marcu, A.; Liang, Y.; Grant, J.R.; Maciejewski, A.; Wishart, D.S. Heatmapper: web-enabled heat mapping for all. Nucleic Acid Res. 2016, 44, W147–W153.

40. Huang, D.W.; Sherman, B.T.; Lempicki, R.A. Systematic and integrative analysis of large gene lists using DAVID bioinformatics resources. Nat. Protoc. 2009, 4, 44–57.

41. Takaya, T.; Kawamura, T.; Morimoto, T.; Ono, K.; Kita, T.; Shimatsu, A.; Hasegawa, K. Identification of p300-targeted acetylated residues in GATA4 during hypertrophic responses in cardiac myocytes. J. BIol. Chem. 2008, 283, 9828–9835.

42. Morimoto, T.; Sunagawa, Y.; Kawamura, T.; Takaya, T.; Wada, H.; Nagasawa, A.; Komeda, M.; Fujita, M.; Shimatsu, A.; Kita, T.; Hasegawa, K. The dietary compound curcumin inhibits p300 histone acetyltransferase activity and prevents heart failure in rats. J. Clin. Invest. 2008, 118, 868–878.

43. Katagiri, T.; Sunagawa, Y.; Maekawa, T.; Funamoto, M.; Shimizu, S.; Shimizu, K.; Katanasaka, Y.; Komiyama, M.; Hawke, P.; Hara, H.; Mori, K.; Hasegawa, K.; Morimoto, T. Ecklonia stolonifera Okamura extract suppresses myocardial infarction-induced left ventricular systolic dysfunction by inhibiting p300-HAT activity. Nutrients 2022, 14, 580.

44. Liu, W.; Brown, K.; Legros, S.; Foley, A.C. Nodal mutant eXtraembryonic ENdoderm (XEN) stem cells upregulate markers for the anterior visceral endoderm and impact the timing of cardiac differentiation in mouse embryoid bodies. Biol. Open 2012, 1, 208–219.

45. Sakurai, H.; Okawa, Y.; Inami, Y.; Nishio, N.; Isobe, K. Paraxial mesodermal progenitors derived from mouse embryonic stem cells contribute to muscle regeneration via differentiation into muscle satellite cells. Stem Cells 2008, 26, 1865–1873.

46. Wang, T.; McDonald, C.; Petrenko, N.B.; Leblanc, M.; Wang, T.; Giguere, V.; Evans, R.M.; Patel, V.V.; Pei, L. Estrogen-related receptor α (ERRα) and ERRγ are essential coordinators of cardiac metabolism and function. Mol. Cell. Biol. 2015, 35, 1281–1298.

